# PhenoEncoder: A Discriminative Embedding Approach to Genomic Data Compression

**DOI:** 10.1101/2024.12.06.625879

**Authors:** Gizem Taş, Eric Postma, Marleen Balvert, Alexander Schönhuth

**Affiliations:** Department of Econometrics and Operations Research, Tilburg University, Warandelaan 2, 5037AB Tilburg, The Netherlands; Department of Cognitive Science and Artificial Intelligence, Tilburg University, Warandelaan 2, 5037AB Tilburg, The Netherlands; Genome Data Science, Faculty of Technology, Bielefeld University, Universitätsstraße 25, 33615 Bielefeld, Germany

**Keywords:** Genomic Feature Extraction, Discriminative Embedding, Autoencoder

## Abstract

Exploring the heritability of complex genetic traits requires methods that can handle the genome’s vast scale and the intricate relationships among genetic markers. Widely accepted association studies overlook non-linear effects (epistasis), prompting the adoption of deep neural networks (DNNs) for their scalability with large genetic datasets and ability to detect complex relationships. However, the curse of dimensionality continues to limit the potential of DNNs, underscoring the critical need for dimensionality reduction for suitably sizing and shaping the genetic inputs, while preserving epistasis.

Linkage disequilibrium (LD), a measure of correlation between genetic loci, offers a pathway to genome compression with minimal information loss. Using LD, the genome can be divided into smaller genomic regions, i.e., haplotype blocks, which can be locally compressed using deep autoencoders. While autoencoders excel at preserving the main non-linear patterns, they still risk losing phenotype-relevant information when dominated by other sources of genetic variation.

We propose a novel approach, PhenoEncoder, that incorporates phenotypic variance directly into compression. This single nucleotide polymorphism (SNP)-based pipeline employs multiple autoencoders, each dedicated to compressing a single haplotype block. The window-based sparsity of the model eases the computational burden of simultaneously processing numerous SNPs. Concurrently, an auxiliary classifier predicts the phenotype from the compressed haplotype blocks. Epistasis is processed both within and between haplotype blocks by maintaining non-linearity in the autoencoders and the classifier. Through joint optimization of the compression and classification losses, PhenoEncoder ensures that disease-causing patterns are highlighted during compression.

Applied to protein expression and simulated complex phenotype datasets, PhenoEncoder demonstrated enhanced generalizability in downstream classification tasks compared to standard autoencoder compression. By enabling phenotype-aware compression, PhenoEncoder emerges as a promising approach for discriminative genomic feature extraction.

## 1 Introduction

The genetic architecture of a disease is considered complex when a combination of several genomic factors contribute to the phenotype [1,2]. Complex dis-eases such as Alzheimer’s disease or amyotrophic lateral sclerosis (ALS) can be driven by the combined impact of numerous genetic variants with small effect sizes [3,4,5], or even by the indirect regulatory influences from the peripheral markers. Moreover, the *omnigenic model* assumes the involvement of virtually all genes in shaping phenotypic outcomes [6], which explains the great challenges one encounters when trying to disentangle the genetic landscape of complex diseases. In particular, many of the effects involved may depend on each other via non-additive relationship, which prevents their easy revelation via standard, genome-wide association study type approaches. The phenomenon expressing such relationships, driving the major difficulties in the discovery of disease-related genetic variants is referred to as *epistasis*, which reflects the existence of causal effects due to non-linear interactions between genetic variants [7,8]. In an overall account, the mechanisms determining the occurrence, heritability, and progression of complex diseases can only be unraveled through a collective analysis of such factors.

The sheer volume of genetic variants often surpasses the number of observations in genomic data [9]. Situated in a severely sparse vector space, the high-dimensional input—possibly featuring multicollinearity or redundancy among variants—gives rise to the *curse of dimensionality* : an issue that undermines the generalizability of predictive models [10]. In view of the immense scale of the genome, transforming the raw genomic data into viable inputs for subsequent computational methods, i.e. *genomic feature extraction*, is an integral part of studying complex disease genomics [11]. This transformation involves encoding the typically high-dimensional genomic data into a lower-dimensional representation, for instance, an embedding, that preserves the task-dependent relevant characteristics of the underlying biology.

A prominent method to produce compact embeddings while preserving non-linearity is through deep autoencoders [12]. While minimizing the discrepancy between the input and the output, deep autoencoders inherently learn salient properties of the data, i.e., the latent features, in the hidden layers, which operate as an information bottleneck by being of lower dimensionality than the input and output [13,14]. Following the advent of technologies yielding high-throughput omics data, the field of bioinformatics has embraced autoencoders for their unmatched potential to automatically derive meaningful insights from this vast data trove. A variety of types of autoencoders, namely vanilla autoen-coders [15], denoising autoencoders [16], and variational autoencoders [17,18,19], have been successfully utilized for representation learning without the need for labels, across diverse forms of genomics data. However, deep autoencoders, with their densely interconnected architectures, are not immune to the *curse of dimensionality* and may still pose computational challenges when attempting to process large genomic data as a single vector input. To alleviate this issue, the genome can be fragmented into, for instance, local windows [20], genes [21], or linkage disequilibrium (LD) regions [22,15], as an effective preprocessing strategy. Building on the known correlations between genetic variants [23], we define the genome segmentation based on its LD structure—an approach previously demonstrated to be efficient by Taş et al. [15].

Unsupervised feature extraction does not always guarantee a downhill connection between the extracted features and phenotypic diversity. A vital yet unresolved challenge is that disease-associated patterns may be obscured by prominent—formally *normal* —sources of genetic variation within the population, including ethnic background, geographic origin, demographic history, and relatedness [24,25,26]. In such cases, the latent features essential for accurately reconstructing genotypes may not necessarily account for the phenotypic variation observed among individuals.

As a potential remedy, one could propose combining unsupervised and supervised learning objectives. Several studies have demonstrated that incorporating unsupervised components into classification tasks not only boosts classification performance and generalization on benchmark datasets but also augments training data by leveraging unlabeled samples within a semi-supervised learning framework [27,28,29,30]. A shared inference from these works is that the abstract representations generated purely through supervised tasks, such as in the hidden layers of neural networks, may remain under-constrained, leading to overfitting and an inability to recognize true characteristics of the data [30]. Integrating both supervised and unsupervised losses helps strike a balance through multi-task learning. In essence, while minimizing the unsupervised loss acts as a regularizer, minimizing the supervised loss steers representation learning toward a latent space that is informative of a specific target. The combined losses offer a symbiotic benefit of regularization and representational relevance.

The aforementioned semi-supervised learning studies commonly regard the unsupervised task as auxiliary, with the primary focus on the supervised task. The reverse perspective, which views supervision as auxiliary, has also been explored in the realm of feature extraction via autoencoders, although this shift is more conceptual than methodological. Dinçer et al. [31] materialize this idea for generating confounder-free embeddings of gene expression profiles by incorporating an adversarial loss, derived from confounder prediction, into the autoen-coder’s loss. Alternatively, the supervised learning objective might not always be embodied in an auxiliary model, yet it can still complement the unsupervised task implicitly. For example, Razakarivony and Jurie [32] substitute the reconstruction error in standard autoencoder training with hinge loss, to constrain the latent space (i.e., “manifold”, as termed by the authors) for reconstructing the positive samples better than the negative ones. Class separability is implied through a spatial modification of the reconstruction targets in the work of Nousi and Tefas [33], such that the autoencoders aim to reconstruct the data points that are now moved closer to their respective class centers, and distant from the rival class centers in the original space. Zhou et al. [34] insert a local Fisher discriminant regularizer, a supervised linear dimensionality reduction technique [35], on the hidden layer activations of the autoencoder to maximize inter-class distances. Regularization of the autoencoders is again harnessed by Hu et al. [36] to impose a label consistency constraint on the neuron activations through penalizing the distance of samples to their class centers. Moreover, a population genetics study by Geleta et al. [20], directly conditions the compression on the ancestry labels using ancestry-specific autoencoders.

Inspired by prior research, we envision that crafting a latent feature space that accentuates phenotypic separability while retaining the intricate relationships within the data should be the key to lifting the veil on the genetic landscape of complex diseases. To achieve this, one must recognize that phenotypic separability may depend on epistatic causal effects. Therefore, the supervised component should account for possible non-additive interactions between variants across the entire genome in a manner compatible with the window-based processing of the genome. Complex interactions may occur both within and between these windows, and inevitably require non-linearity for their recognition. This can be accomplished through an auxiliary fully connected neural network trained on the supervised learning objective on top of autoencoder compression. In light of the above, we present the PhenoEncoder: a multi-autoencoder framework designed to produce discriminative embeddings. PhenoEncoder compresses high-dimensional genomic input into a lower-dimensional unified latent feature space, by concatenating multiple information bottlenecks, while also preserving discriminatory phenotypic information—–disease status, for example–—through the auxiliary classifier. Customized to tackle the specific challenges encountered in complex disease genomics, the PhenoEncoder provides four key contributions: (1) scalability to large genome data, in particular single nucleotide polymorphism (SNP) data, within the limits of computational feasibility, while (2) maintaining non-additive effects both within and between the fragments of input, (3) an epistasis-enabled compression suitable for complex phenotypes, and, most importantly, (4) the potential to highlight the pathological patterns in the latent space even when they are non-dominant.

## 2 Methods

This section outlines the data preprocessing, the PhenoEncoder model, its training strategy, and evaluation protocols. First, we detail the window-based pre-processing of the genome. Next, we introduce the PhenoEncoder architecture, loss function, training algorithm, followed by its evaluation in comparison with a standard autoencoder. Finally, we describe the two datasets and their respective use cases in this study.

### 2.1 Window-based Preprocessing of the Genome

To efficiently manage the high dimensionality of SNP data, we follow the same approach as in Taş et al. [15] by segmenting the genome into manageable units based on patterns of LD, where certain alleles are inherited together more often than by chance [37]. These segments, known as haplotype blocks, represent regions of the chromosome that contain only a few distinct haplotypes, or sets of alleles passed down from one parent [38]. By focusing on regions with limited historical recombination, we efficiently compress the data while preserving the underlying genetic correlations. To define the haplotype block boundaries, we use the implementation provided by PLINK 1.9 [39]. To create large, densely packed blocks, which are ideal for dimensionality reduction, we modified the default parameters as follows: the confidence interval for strong LD was adjusted to range between 0.5 and 0.85, the upper recombination threshold was reduced from 0.9 to 0.7, and the SNP window size was increased from 200 Kb to 10 Mb.

### 2.2 The PhenoEncoder Model

#### Architecture

We designed a modular architecture for the PhenoEncoder, as depicted in Figure 1, using autoencoders along with an auxiliary classifier as fundamental building blocks. In this configuration, the autoencoders process multiple input vectors -the haplotype blocks -in parallel to produce multiple reconstructed output layers. The latter use a custom activation function formu-lated by Taş et al. [15] for recoded SNP values of ordinal nature, as defined in Equation S3 in Supplementary Note 1. The input layer of the classifier is then formed by concatenating the bottleneck layers side by side.

**Fig. 1:**
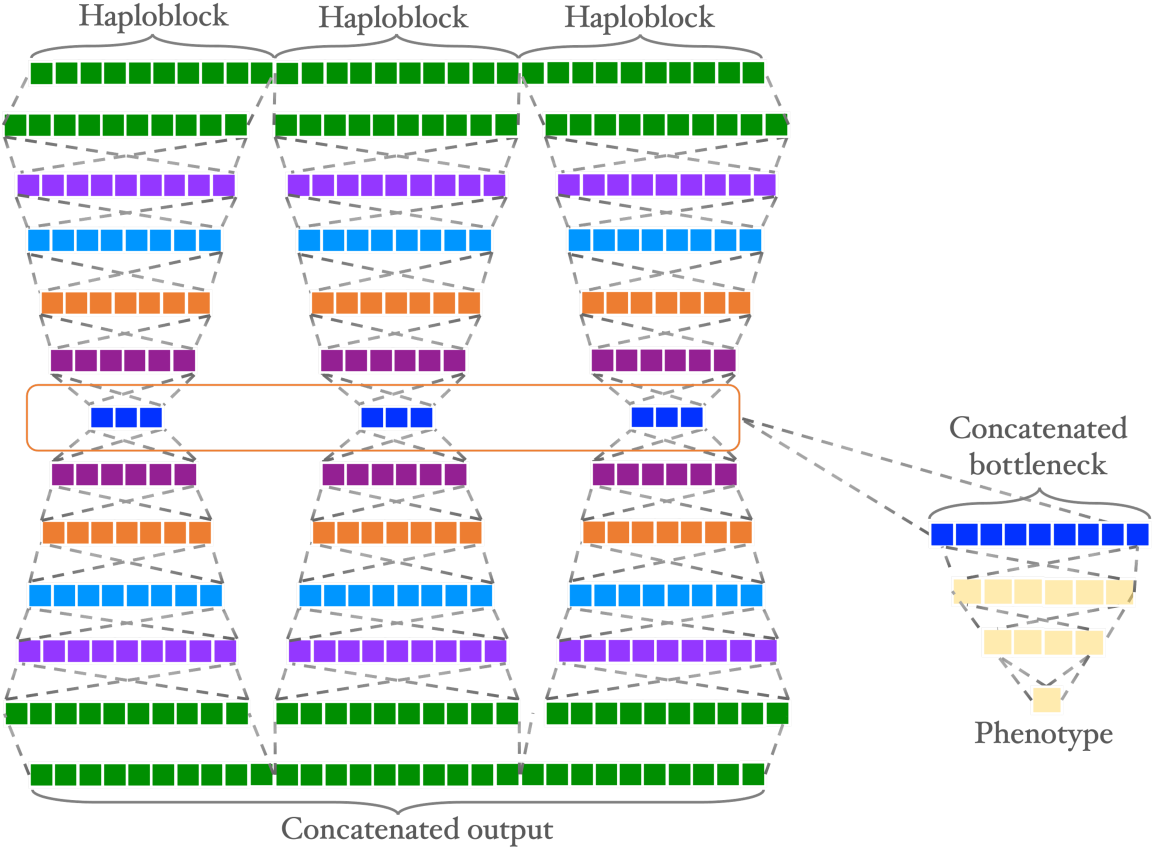
Architecture of the PhenoEncoder. The PhenoEncoder consists of multiple autoencoders that each compress a fragment of the data, where the input data is segmented into haplotype blocks. The concatenation of the autoencoders’ compressions of the haploblocks forms the input to the classifier, which predicts a sample’s phenotype. By combining the loss of the autoencoders with the loss of the classifier -see Equation 1 for details -the data compression of the autoencoders is steered towards retaining information that is relevant for phenotype classification. The autoencoder loss is based on the concatenation of the output layers of the individual autoencoders to ensure that the loss is independent of the number of autoencoders used.

Each autoencoder in Figure 1 consists of an *encoder* which progressively compresses the input, a *bottleneck layer* where the abstract representation is achieved, and a *decoder* which aims to reconstruct the input from the bottleneck layer. This way, the autoencoders prove useful for feature extraction.

Overall, the autoencoders compress their input in a self-supervised manner, capturing meaningful data patterns in a lower-dimensional abstract representation. When combined with the auxiliary classifier, the compression becomes supervised by incorporating phenotype information, resulting in a discriminative embedding in the latent space.

We implemented a fully-connected autoencoder architecture adaptable to various input sizes formed by *shape* to compatibly adjust the width of the layers, *number of hidden layers* to set the depth of the network, and *bottleneck dimension* to determine the latent space dimensionality. We built the autoen-coders using the resulting optimal strategy following Taş et al. [15], compatible with diverse haplotype blocks. More details concerning the autoencoder implementation and the selected hyperparameters for this strategy are provided in Supplementary Note 1.

The auxiliary classifier takes the concatenated bottleneck outputs as input and learns to classify case and control samples. The classifier architecture is optimized through a grid search concerning the depth (number of hidden layers) along with the width (the number of nodes per layer) of the sub-network, the dropout rate, the learning rate, and the batch size, see Table S1 of Supplementary Note 2. The optimized network consists of two fully-connected hidden layers, both with Leaky Rectified Linear Unit (Leaky ReLU) transfer functions [40], initialized using the He uniform variance scaling initializer [41], and regularized with a dropout layer [42] in between (for optimized dropout rates, see Table S1 of Supplementary Note 2). Finally, the 1-dimensional output layer is equipped with a sigmoid activation function, tailored for binary classification.

#### The Joint Loss

The PhenoEncoder model aims to minimize a joint loss func-tion *L*_total_:

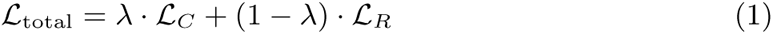

where *L_C_*is the classification loss, which guides the latent space to capture features that are predictive of the target phenotype, while *L_R_* is the reconstruction loss that ensures that the embedding accurately represents the underlying data patterns and provides regularization for the supervised tasks [30].

The parameter λ *ɛ* 2 (0, 1) is the loss weight coefficient that controls the balance between these two tasks and must be carefully tuned to manage the trade-off between overfitting in supervised learning and an under-constrained (i.e. phenotype-agnostic) representation learning. We provide further insight into the impact of λ on balancing two tasks in Supplementary Figures S2, S3 and and include the details of λ tuning experiments in Supplementary Note 3.

#### Alternating Training Algorithm

PhenoEncoder training begins with the respective pre-training of its two sub-networks: the autoencoders and the auxiliary classifier. First, the autoencoders are pre-trained to minimize the reconstruction loss (*L_R_*), measured by mean squared error (MSE). To compute the MSE, the outputs of multiple autoencoders are concatenated into a single vector on the same scale as the original input, allowing for the calculation of a single MSE value regardless of the number of autoencoders in the system (see Figure 1). Then, using the data compression obtained by the pre-trained autoencoders, the classifier is pre-trained to optimize the binary cross-entropy (BCE) loss (*L_C_*) between the true phenotype labels and the sigmoid output.

Once both sub-networks are initialized, we implement an alternating training procedure in which the PhenoEncoder and the auxiliary classifier weights are optimized in turns. In one epoch, we freeze the classifier’s weights and train the PhenoEncoder to minimize the joint loss function *L*_total_ in Equation 1. This way, only the gradients pertaining to the autoencoders become effective. In the subsequent epoch, we freeze the autoencoders while the classifier weights are unfrozen and updated to minimize the BCE loss (*L_C_*).

This process is summarized in Algorithm 1, where *N_ae_* and *N_c_* are the number of pre-training epochs for the autoencoder and the classifier, respectively, and *N_pe_* is the number of joint training epochs for the PhenoEncoder. We tuned *N_ae_* and *N_c_* to the point where the losses just begin to converge. All training steps utilize the Adam optimizer [43] with an initial learning rate of 0.0001. The optimal hyperparameter combinations for the PhenoEncoder configuration are presented in Table of Supplementary Note 2.

|  |  |
| --- | --- |
| `1: Pre-train Autoencoders for $N_{ae}$ epochs 2: Pre-train Classifier for $N_c$ epochs 3: for $i = 1$ to $N_{pe}$ do if $i$ is even then Freeze Classifier Train PhenoEncoder for 1 epoch else Freeze Autoencoders Unfreeze Classifier Train Classifier for 1 epoch end if end for` | where<br>$N_{ae}$: The number of Autoencoder pre-training epochs.<br>$N_c$: The number of Classifier pre-training epochs.<br>$N_{pe}$: The number of PhenoEncoder joint training epochs. |

#### Performance Evaluation

The PhenoEncoder is primarily designed to obtain discriminative embeddings from large genotyping data, which support the generalizability of downstream classification tasks. The classification performance of the PhenoEncoder is assessed using five metrics derived from the auxiliary classifier’s output: accuracy, precision, recall, the areas under the receiver operating characteristic (AUC-ROC) and precision-recall (AUC-PRC) curves.

Secondarily, the quality of the compression is evaluated using MSE alongside the percentage of correctly predicted SNPs by comparing the original and reconstructed SNP vectors, a metric called *SNP reconstruction accuracy*, as an indicator of information loss for the autoencoder compression*^†^*.

#### Comparison of Autoencoder and PhenoEncoder Compression

The performance of the PhenoEncoder was ultimately evaluated by comparing its discriminative power on unseen test data to a baseline model. As the aim of the PhenoEncoder is to incorporate phenotype-specific information in its embeddings, we compare the PhenoEncoder to classical (multiple) autoencoders which correspond to the autoencoder component within the PhenoEncoder, and are trained only to minimize the reconstruction loss, ceteris paribus*^‡^*. Both models create an embedding of the data, where we hypothesize that a classification based on the PhenoEncoder embedding outperforms a classification based on the classical autoencoder embedding.

Recalling that the ultimate goal of our methodology is to create embeddings that support inductive learning, we eventually evaluate the worth of the PhenoEncoder through the performances of two downstream classifiers—logistic regression and a multi-layer perceptron (MLP)—on the generated embeddings. The downstream MLP differs functionally from the PhenoEncoder’s auxiliary classifier, as it is trained separately on the final embeddings after the PhenoEncoder’s training is complete. Both classifiers are assessed using five criteria: accuracy, precision, recall, the areas under the receiver operating characteristic (AUC-ROC) and precision-recall (AUC-PRC) curves. Furthermore, as the PhenoEncoder balances reconstruction loss versus classification accuracy, it likely yields a higher reconstruction error than the autoencoder, which we test below.

### 2.3 Datasets and Use Cases

Our experimental setup for evaluating the PhenoEncoder framework consists of two distinct pipelines on two datasets, as illustrated in Figure 2. The first pipeline depicted in the upper branch leverages a small, manageable dataset, merely to validate the PhenoEncoder, providing an initial test of its ability to extract meaningful discriminative embeddings for a relatively simple phenotype classification task. With this pipeline, we present our preliminary results. The second pipeline in the lower branch describes our complete methodology tested on the VariantSpark Synthetic Data, a high-dimensional SNP dataset from the 1000-Genomes Project [44], with complex phenotypes generated by simulated epistatic interactions, leading to the principal findings in this study.

**Fig. 2:**
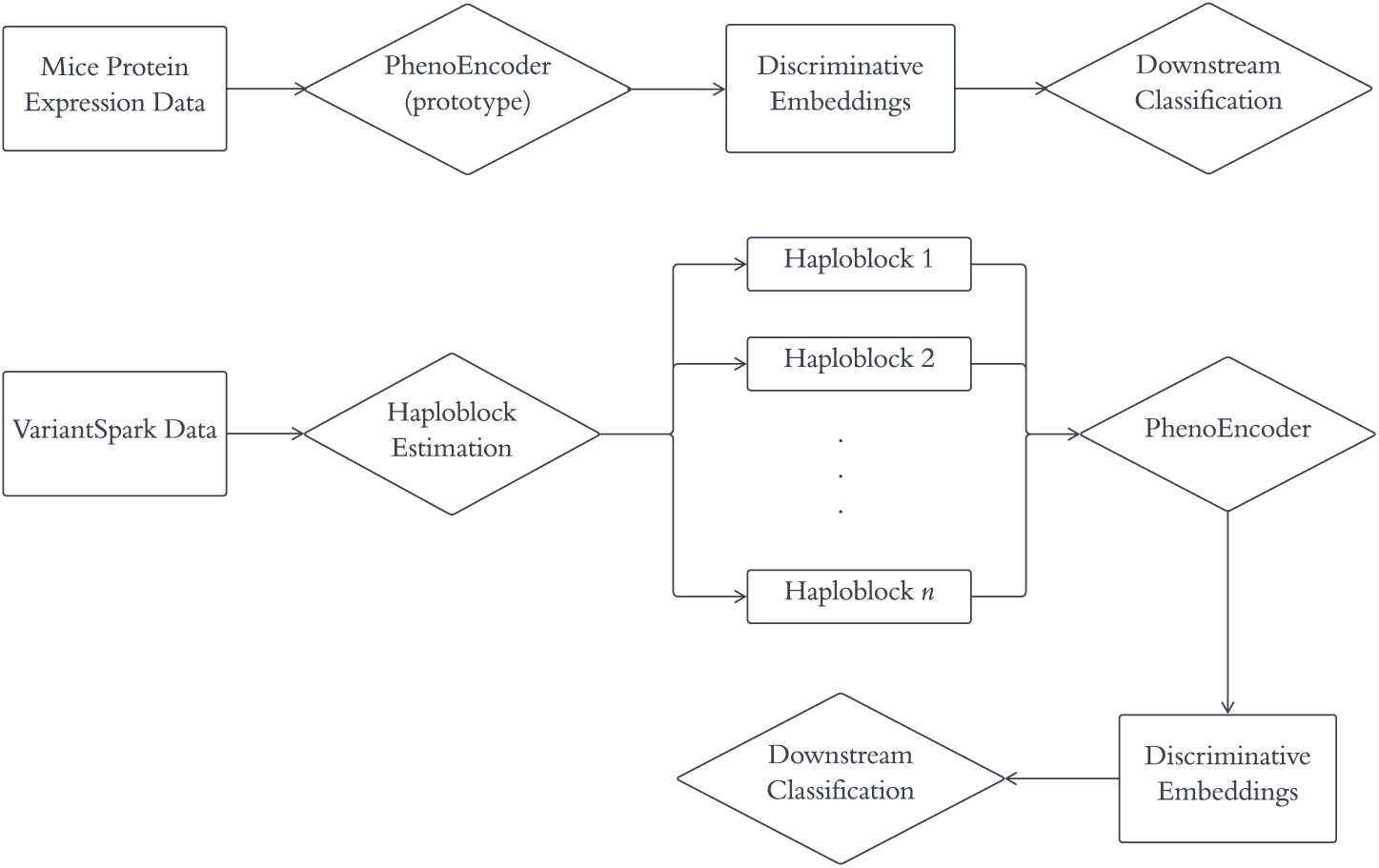
Overview of the experimental setup. We conduct two experiments: the first uses a small dataset and aims to validate the PhenoEncoder (top flow chart), and the second is our principal experiment using a complex SNP dataset (bottom flow chart).

#### First Test Pipeline: Mice Protein Expression Data

In order to test whether the rationale behind the PhenoEncoder works, we use a very small dataset on mice, where the considered phenotype is Down Syndrome. The size of the data, containing only 77 features, allows us to conduct a thorough analysis on results obtained by the PhenoEncoder compared to baseline models. Although the relatively small dimensionality of this particular dataset does not necessarily present the computational challenges associated with complex genetic traits, the multiplex assortment of mice in this instance provides an ideal sample for testing a prototype of the PhenoEncoder model. We utilize this sample primarily to validate the effectiveness of the PhenoEncoder in capturing and highlighting phenotypic variations of interest (control or trisomic) within the latent space. Given the manageable size of the original feature vector, the input is processed as one piece through a single autoencoder which produces a linear-activated reconstructed output, and the classifier processes the autoencoder’s bottleneck into one sigmoid-activated output node, see Figure 3.

**Fig. 3:**
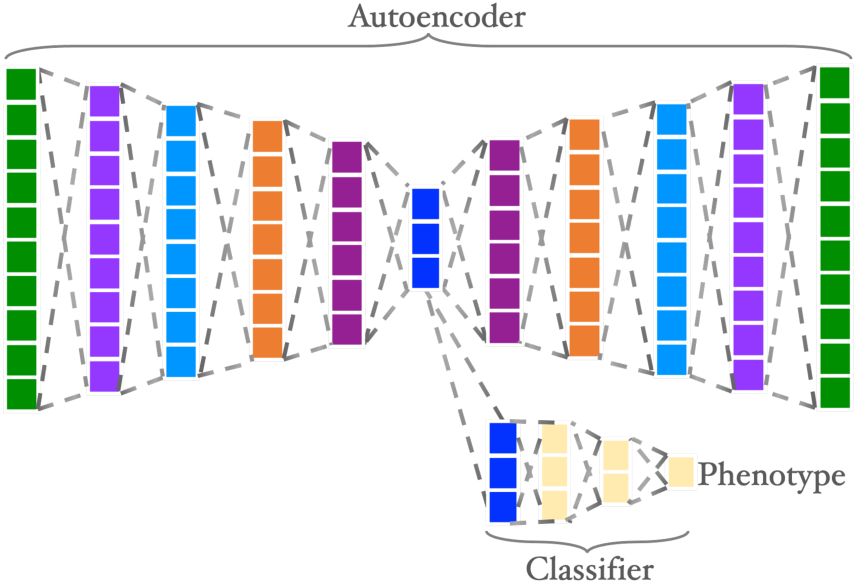
Architecture of the prototype model. Optimal hyperparameters are provided in Supplementary Note 5.

The multivariate dataset features continuous expression levels of 77 proteins from the cerebral cortex of mice in 1080 measurements, each corresponding to an independent mouse sample. 570 of these mice belong to the control group, while the remaining 510 are labeled ‘trisomic’ based on their Down Syndrome status. The mice can also be grouped by their behavior (either stimulated to learn or not), and their treatment status (memantine drug or saline injection). The dataset is obtained from the public through the UCI Machine Learning Repository [45].

#### Second, Full Pipeline: VariantSpark Synthetic Data

The publicly avail-able dataset from VariantSpark [46] consists of real genotypes (over 81 million genetic variants) of 2,504 samples as part of the 1000-Genomes Project [44] for which binary phenotype labels are simulated using the Polygenic Epistatic Phenotype Simulator (PEPS) [47]. Among those, we selected a complex simulated phenotype which is associated with individual effects as well as epistatic—both pairwise and higher-order—interactions between a subset of 253 randomly selected variants, known as the ‘truth variants’.

##### Initial feature selection

Following the estimation of haplotype block boundaries as prescribed by the divide-and-compress approach by Taş et al. [15] to high-dimensional SNP input, the number of haplotype blocks can still amount to thousands, which would lead to computational challenges. Therefore we decided to benefit from the ground truth SNPs available in this particular dataset, for an initial feature selection.

We identified the ‘truth blocks’ that contain at least one truth variant. The random initial selection of the truth variants and limited size of the sample may result in a number of isolated variants remaining off-blocks. A truth variant not covered by any block is then merged with the nearest block. Thus, we obtained 253 haplotype blocks which contain a total of 6,658 variants including 253 truth variants, corresponding to 100% of the true causal effects in this instance.

#### Data Splitting

To begin with, we stratified the Mice Protein Expression dataset by three available labels (phenotype, behavior and treatment) and reserved 20% for testing, so that all training and hyperparameter tuning experiments were performed exclusively on the remaining 80% of the data. The testing subset was only utilized for reporting the final results. As for the VariantSpark samples, 10% of the samples were allocated for testing as we anticipated a need for a larger training portion due to the higher dimensionality of the dataset and the heightened risk of overfitting.

## 3 Results

This section presents the outcomes of the two pipelines illustrated in Figure 2. Preliminary results from the first pipeline were obtained using the prototype model on the small-scale Mice Protein Expression Data, while principal findings from the second pipeline were derived from the VariantSpark Synthetic Data. All computations were performed on a system featuring an Intel Xeon Gold 6242R processor clocked at 3.10 GHz, 376 GB of RAM, and running Rocky Linux version 8, provided by the high-performance computing (HPC) facilities of the Utrecht Bioinformatics Center (UBC).

### 3.1 Preliminary Findings

The prototype model of the PhenoEncoder based on the mouse gene expression data allowed us a glimpse into the potential value of the methodology through an inexpensive look at the intended study outcomes.

#### Prototype Model Classification Performance

The optimal prototype model (see Supplementary Note 5 for details on the hyperparameter tuning) configured with λ = 0.7 (see Table S2 of Supplementary Note 3 for the details regarding the tuning of λ) was trained once on the entire training subset, including the validation samples. Evaluation of this model on the unseen test data demonstrated its remarkable ability to distinguish trisomic mice from control samples, with all classification metrics in Table 1 exceeding 99%.

**Table 1:** Test evaluation results for the prototype model.

| $\lambda$ | Accuracy | Precision | Recall | AUC-ROC | AUC-PRC |
| --- | --- | --- | --- | --- | --- |
| 0.7 | 0.9954 | 1.0000 | 0.9902 | 0.9940 | 0.9962 |

#### Comparison to the Baseline Embeddings

The bottleneck representations obtained from the prototype model constitute the PhenoEncoder embeddings, which can be compared with the classical autoencoder embeddings*^§^* through visualizing the latent spaces. The visualization serves as a tool for interpreting and judging the effect of our approach compared to the baseline unsupervised compression merely using an autoencoder. We applied Uniform Manifold Approximation and Projection (UMAP) [48] to the resulting embeddings, as shown in Figures 4 and 5, to reveal the underlying structure in a 2-dimensional space. Visualization of the data compressed by an unsupervised autoencoder gives Figure 4, where the data points are colored by behavior (left figure) and phenotype (right figure). Recall that we aim to obtain a data representation that allows us to separate mice with Down Syndrome from healthy controls. Figure 4 shows that the current data compression does not facilitate an easy classifica-tion by phenotype, while the two clusters based on behavior are visible in the latent representation, which may be the dominant source of variation, but is not our main interest. On the other hand, when using the simplified PhenoEn-coder prototype, the latent space does lead to a proper separation of the mice by phenotype, as shown in Figure 5.

**Fig. 4:**
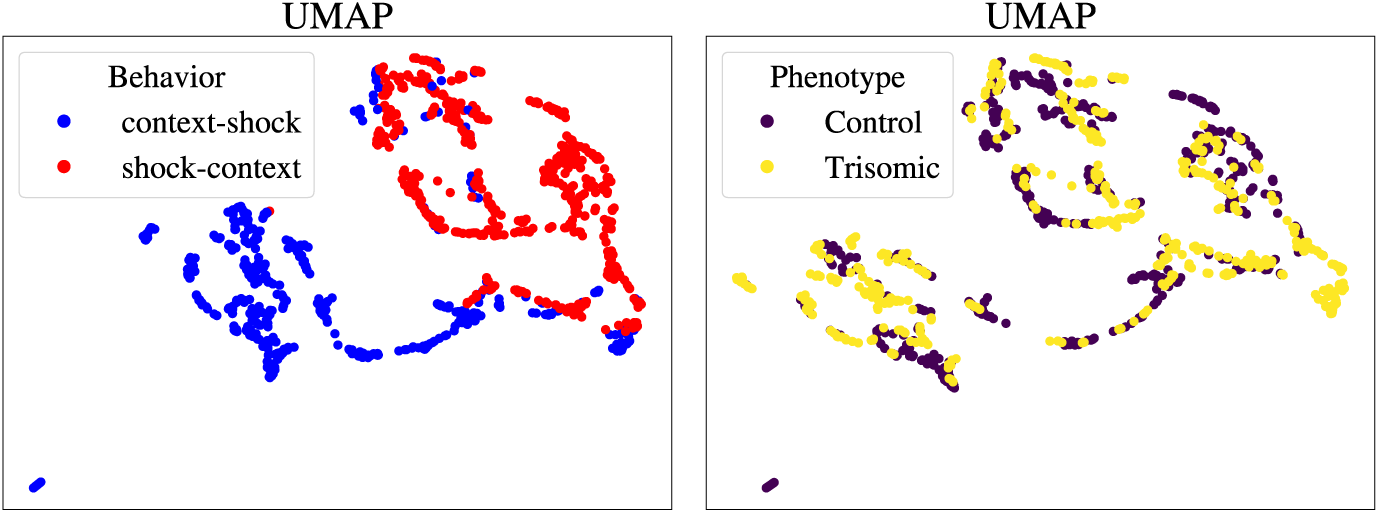
Visualizations of the Mice Protein Expression Dataset embedded by the autoencoder. The unsupervised compression generates a latent space encoded by the main factors of variation in the data: the behavior. On the left, we observe a clear separation between behavior classes. However, the phenotypic variation of interest seems less evident in the resulting latent space as shown by the lack of distinction between trisomic and control samples on the right.

**Fig. 5:**
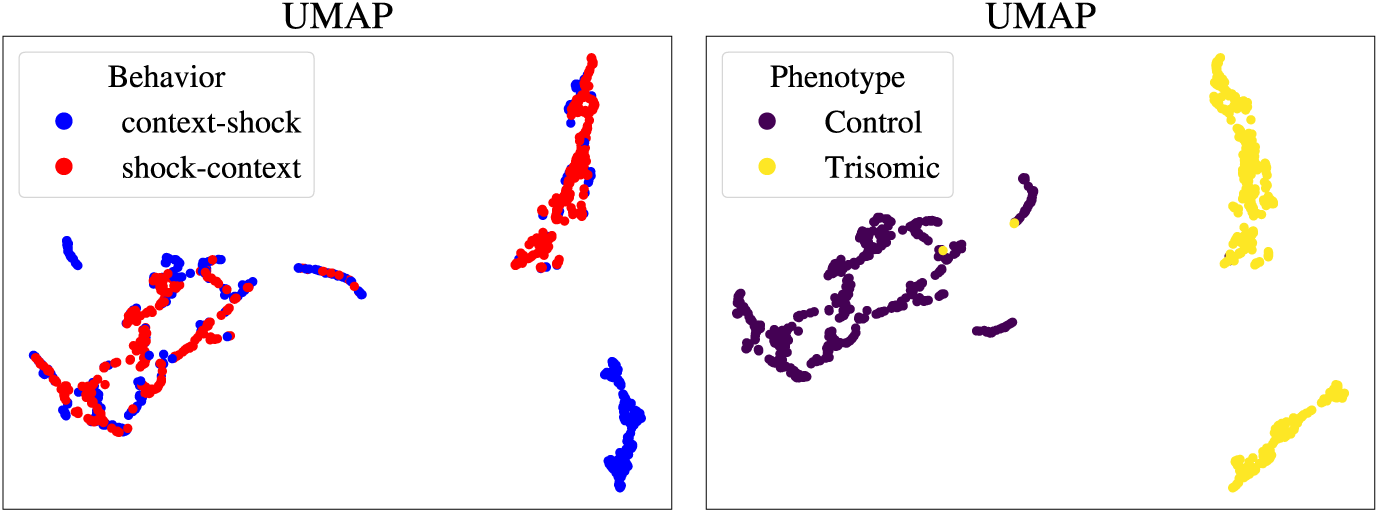
Visualizations of the Mice Protein Expression Dataset embedded by the PhenoEncoder prototype. Our supervised compression approach manages to prioritize the phenotype-relevant features over the dominant substructures in the original data. The visualization on the left hints that the latent space produced by the PhenoEncoder is not primarily driven by the behavior-related features anymore, but rather by the phenotype information. When embedded by the PhenoEncoder, the separation between case and control groups on the right is notably more visible than the autoencoder vi-sualizations displayed above.

The improved separability of the phenotype status in the contrasted latent spaces in Figures 4 and 5 is corroborated by the subsequent classification experiments. Table 2 shows that both linear (logistic regression) and non-linear (MLP) classification methods demonstrate higher discriminative ability when performed on the latent space embedded by the PhenoEncoder compared to the latent space obtained by the autoencoder. When succeeded by logistic regression, the PhenoEncoder’s compression leads to a roughly 19% increase in test accuracy and 20% in test AUC-ROC, whereas the improvement is even more apparent for the MLP classifier by around 24% and 27% respectively.

**Table 2:** Preliminary comparison of autoencoder (AE) and PhenoEncoder (PE) embeddings of test samples through evaluation of downstream classification performance.

| Method | Model | Accuracy | Precision | Recall | AUC-ROC | AUC-PRC |
| --- | --- | --- | --- | --- | --- | --- |
| Logistic Regression | AE | 0.5833 | 0.5938 | 0.3725 | 0.5722 | 0.5175 |
|  | PE | **0.7731** | **0.7154** | **0.8627** | **0.7779** | **0.6821** |
| Multi-layer Perceptron (MLP)* | AE | 0.5509 | 0.5229 | 0.5588 | 0.6018 | 0.5734 |
|  | PE | **0.7917** | **0.8701** | **0.6569** | **0.8779** | **0.8290** |

\* The MLP architecture here mirrors the classifier sub-network within the prototype model and is trained for 25 epochs using a learning rate of 0.0001 and a batch size of 32.

**Table 3:** Test evaluation results for the PhenoEncoder.

| $\lambda$ | SNP rec. acc. | Accuracy | Precision | Recall | AUC-ROC | AUC-PRC |
| --- | --- | --- | --- | --- | --- | --- |
| 0.3 | 0.8744 | 0.6295 | 0.6204 | 0.6746 | 0.6420 | 0.6339 |
| 0.7 | 0.8583 | 0.5578 | 0.5581 | 0.5714 | 0.6033 | 0.5907 |

Our preliminary findings validate the positive impact of integrating phenotype information into compression through the PhenoEncoder model. In this initial step, the latent features captured by the simplified model proved more phenotype-relevant than those of the autoencoder, an outcome endorsed by the classification and visualization experiments. Therefore we believe that the full PhenoEncoder model, with its scalable multiple autoencoder architecture, is worth pursuing to produce genotype embeddings better suited for separability with regard to complex traits.

### 3.2 Principal Findings

We trained the full PhenoEncoder model on the VariantSpark data, to provide outcomes particularly within the context of complex disease genomics. Details of the computational resources utilized for these experiments are listed in Supplementary Table S4. Details regarding hyperparameter tuning can be found in Supplementary Note 5.

#### PhenoEncoder Classification Performance

We constructed two distinct PhenoEncoder models using the hyperparameter setting in Table S1, one to be compiled with λ = 0.3 and the other with λ = 0.7, as indicated by λ tuning results in Table S3 of Supplementary Note 3. Both models were trained on the entire training subset and evaluated on the so far unseen test subset. Table 3 indicates better overall quality of the PhenoEncoder when compiled with λ= 0.3, reflected in higher accuracy (0.6295), AUC-ROC (0.6420), and other classification metrics as well as better SNP reconstruction accuracy (0.8744).

#### Comparison to the Baseline Embeddings

The main goal of the PhenoEncoder is to steer data compression towards capturing phenotype-related genomic characteristics. To test whether this was achieved, we apply two classifiers, namely logistic regression and MLP, to classify cases versus controls from the data compressed by the autoencoder and from the data compressed by the PhenoEncoder. As displayed in Table 4, logistic regression applied to the PhenoEncoder embeddings of test samples outperforms classification based on the autoencoder compression ¶ by approximately 5% in both accuracy and AUC-ROC for λ = 0.7, and by 9% for λ = 0.3. The improvement is even more pronounced with the MLP model, where the PhenoEncoder compiled with λ = 0.3 results in over a 10% boost in accuracy and a 9% gain in AUC-ROC. Additionally, an improvement of nearly 11% in AUC-ROC is observed with λ = 0.7.

**Table 4:** Comparison of autoencoder (AE) and PhenoEncoder (PE) embeddings of test samples through evaluation of downstream classification performances.

| Method | Model | Accuracy | Precision | Recall | AUC-ROC | AUC-PRC |
| --- | --- | --- | --- | --- | --- | --- |
| Logistic Regression | AE | 0.5378 | 0.5385 | 0.5556 | 0.5378 | 0.5223 |
| | PE ($\lambda : 0.3$) | **0.6215** | **0.6183** | **0.6429** | **0.6214** | **0.5768** |
| | PE ($\lambda : 0.7$) | 0.5936 | 0.5909 | 0.6190 | 0.5935 | 0.5570 |
| Multi-layer Perceptron (MLP)<sup>*</sup> | AE | 0.5139 | 0.5526 | 0.1667 | 0.5259 | 0.5371 |
| | PE ($\lambda : 0.3$) | **0.6175** | **0.6056** | **0.6825** | 0.6163 | 0.5732 |
| | PE ($\lambda : 0.7$) | 0.5817 | 0.5882 | 0.5556 | **0.6330** | **0.6026** |

\* The MLP here shares the same architecture as the classifier component of the full PhenoEncoder and undergoes 50 epochs of training with a learning rate of 0.0001 and a batch size of 32.

## 4 Discussion

This work introduces a discriminative embedding approach by which to project genotype data into latent spaces. The resulting embeddings can be subsequently used in classification schemes that require to reduce the immense dimensionality of genotype data—note that genotype profiles correspond to vectors of up to 5 million entries. Prior approaches focused on generating embeddings from which to reestablish the original data without notable losses during the reconstruction. However, they were not able to integrate labels, such as for distinguishing between diseased and healthy genotypes, as prevalent in GWAS cohorts, into the compression schemes. Here, we suggest the PhenoEncoder, as a novel approach to consider the labels that accompany the genotypes during the embedding process. Unlike prior approaches to compressing genotype data, PhenoEncoder not only warrants accurate reconstruction of genotypes, but also ensures that embeddings are sufficiently discriminative with respect to their labels.

For this purpose, PhenoEncoder builds on autoencoders at its core, adhering to prior, approved approaches. In addition, PhenoEncoder integrates an MLP, as a classifier that steers the generation of embeddings towards being discriminative. As per its design, the resulting loss function not only penalizes inaccurate reconstruction, but also insufficient classification of the embeddings. In other words, the embeddings already have to carry information about the additional labels; if not, the auxiliary MLP based classifier would be unable to distinguish between the different classes sufficiently accurately.

In our experiments, we assess how effectively two unified latent spaces—one from purely unsupervised compression with multiple autoencoders (baseline) and the other from the PhenoEncoder—retain key data structures essential for accurate classification. Our results demonstrate that PhenoEncoder based genotype embeddings outperform the baseline (by ≈ 10%) when used in subsequent classification tasks. In case of unsupervised feature engineering, the latent space may fail to capture causal relationships between original features, which, lost in compression, will not be recognized by subsequent models. Concerning the heritability of complex traits, this translates to a loss of causal patterns early in the workflow. Therefore, the higher test scores achieved by the PhenoEncoder empirically signify better conservation of class separability in the latent space.

Furthermore, the PhenoEncoder single-handedly achieves the highest test classification accuracy (≈ 63%) and AUC-ROC (≈ 64%), versus downstream logistic regression and MLP models, even by a narrow margin. This outcome may allude to a positive impact of the unsupervised component on the classification performance, thereby verifying the reciprocal relationship between two tasks. As anticipated by Lei et al. [30], the reconstruction loss too has a regularizing effect on the otherwise under-constrained classification loss, enhancing the robustness of the model against overfitting. This is apparent from our results, where both train and test performance are enhanced by the PhenoEncoder. Thus, the concurrent optimization of multiple objectives becomes cooperative.

By adjusting how the data is described in the latent space, the PhenoEncoder produces discriminative embeddings which take the case-control status of samples into consideration, even if the original main variation in the population is driven by phenotype-agnostic factors. Although supervising a deep autoencoder compression is not an unprecedented attempt, the PhenoEncoder brings a novelty to the realm of genomic embeddings by specifically addressing the question of epistasis encountered in complex traits. This novelty can be attributed to two methodological aspects. First, the parallel compression of distinct genomic regions through multiple autoencoders ensures scalability to large data dimensions. Using biological priors such as LD to define the connectivity between layers, we localize and sparsify the architecture such that each region-specific autoencoder focuses on a manageable subset of the genome. Aside from computational efficiency, this structure also provides modularity to the model, a useful feature for adding and removing blocks, as well as for model optimization. Second, granted that each autoencoder was independently supervised by an auxiliary classifier trained on the corresponding limited genomic region, these classifiers would fall short of grasping the epistatic effects between different genomic regions, which may therefore hardly be highlighted in the latent features besides the dominant variation. For the PhenoEncoder however, the supervision over the compression process accounts for potential effects—individual or inter-acting—across the genome owing to the auxiliary classifier being trained on the combination of regional embeddings. Meanwhile, non-linearity is preserved both within and between the blocks via fully-connected neural networks.

In this study we tested the performance of the PhenoEncoder using genotype data from the 1000-Genomes project [44] with a simulated phenotype. As a result, the SNPs that are truly causal of the phenotype are known, allowing us to preselect the genomic regions containing these SNPs and thus enabling a proper testing of the PhenoEncoder without the results being influenced by our pre-selection method. Pre-selection of genomic regions remains necessary, as even though the PhenoEncoder allows for handling high dimensional data, the sheer size of SNP data, containing many millions or even billions of SNPs, still surpasses the dimensions that classification models can handle. In the case of real phenotype data, several pre-selection methods for genomic regions are available, for example one could use biological prior knowledge or select regions based on a GWAS using a very mild *p*-value threshold.

We should herein remark that the supervision signal from the phenotype labels prompts the PhenoEncoder to optimize for classification directly. Accordingly, the discriminative features learned by the PhenoEncoder, albeit customized for separating the classes, do not necessarily lead to larger inter-class distances or smaller intra-class spread, as the model might learn features that enhance decision boundaries rather than maximizing spatial separation. In other words, while a better performance can be indicative of improved decision boundaries for the classifiers, it does not precisely translate to distance-based separability of the samples in the latent space. A classifier’s decision-making logic might rely on complex, non-linear combinations of the learned features and is not analogous to, for example, distance-based clustering algorithms. When the topology of the resulting embeddings is particularly important for downstream analysis, i.e. for clustering, the auxiliary task ought to leverage spatial similarity.

The outcome of the cooperation between the two objectives is by all means contingent upon the delicate balance between the tasks, quantified by the loss weight coefficient λ. Although λ symbolizes the weight of the classification loss in our formulation (see Equation 1), its magnitude has proven not directly proportional to the classification performance, by the suboptimal results with the highest experimented value of 0.9 in Table S3. Unduly weighing the supervised loss may have compromised the intended synergy by either diluting the regularizing effect, which may have induced overfitting, or indirectly obstructing the positive impact from the reconstruction loss, which gradually increases (worsens) as its weight (1 -λ) scales down. In fact, this balance rests on the interdependence of hyperparameters such as the number of joint or pre-training epochs, the learning rate, the batch size, and naturally, λ. In cases where an optimal configuration is hard to achieve, potential users could consider extending the PhenoEncoder’s joint loss by automatic or dynamic loss weighing strategies [49,50,51].

Our findings demonstrate the empirical advantage of the PhenoEncoder embeddings over the baseline with regard to better class separability. Nevertheless, there are still areas that could be refined further by future research. First, as much effort has been put into tuning hyperparameters of the method, the classification component still remains prone to overfitting with increasing data dimensionality. The PhenoEncoder is a multi-input multi-output model. During joint training of the model, although the weights of the auxiliary classifier are frozen, the parallel encoders are still partially optimized to minimize classification loss. This means that as the number of autoencoders increases, which is probable for complex disease research, the complexity of the classifier—as extended by the encoders—will inherently rise, which would exacerbate the vulnerability of the model to overfitting. Except for the previously mentioned smart loss weighing methods, introducing a classifier-free auxiliary supervision through for example similarity learning techniques could amend this issue. But, learning similarities could focus on the geometry of the latent spaces, unlike classification. Also, in order not to lose the edge in maintaining non-linearity between blocks, the low-level embeddings could be fused into a more integrated representation before computing the distance metrics. Another critical aspect is the need to account for confounding factors, which can distort the true relationship between the variables of interest. When confounding might be a concern, to prevent spurious associations, such factors need to be controlled for either by being removed through adversarial learning [31,52], or when available, accounted for as inputs in the model [53]. Lastly, learning the phenotypic similarity between samples rather than point prediction, when implemented properly, could also help indirectly mitigate the issue of confounding.

## 5 Conclusions and Future Work

In this article, we introduced the PhenoEncoder, a novel approach for generating discriminative embeddings from genomic data through a combination of parallel autoencoders and an auxiliary classifier. The PhenoEncoder operates on multiple haplotype blocks in parallel, a way of localizing the compression and sparsifying the complexity of the DNNs working on large genome datasets. In the meantime, the non-linearity is preserved both within individual blocks and across the genomic scale. By integrating phenotype information into the compression process, the PhenoEncoder produces feature vectors that emphasize causal patterns relevant to the phenotype of interest. Our experiments showed that the PhenoEncoder improves downstream classification performance compared to unsupervised compression, with notable gains in both logistic regression and MLP accuracy. These results highlight the value of embedding a discriminative objective into data compression to better capture causal patterns and maintain class separability in the latent space.

Our phenotype-driven compression approach opens new avenues for research, among which comes unraveling the black box, that is, the interpretability of the predictive DNNs. Through the analysis of feature interactions and importance in DNN layers [54,55,56], one can investigate the transformation of the latent space at the haplotype block resolution, identify the genomic regions that contribute most to this transformation, and examine how these contributions relate to the phenotype of interest. This opens the door to uncovering biologically meaningful patterns within compressed genomic data.

## Availability and Implementation

Datasets are available for public use at https://archive.ics.uci.edu/ml/datasets/Mice+Protein+Expression and http://gigadb.org/dataset/100759. Code is available at https://github.com/gizem-tas/phenoencoder.

## Acknowledgments

The authors express gratitude to the High-Performance Compute (HPC) facility, part of the Utrecht Bioinformatics Center (UBC), for providing the essential computing resources for this study.

This work is funded by the Dutch ALS Foundation (Project AV20190010) and the Dutch Research Council (NWO) Veni grant VI.Veni.192.043. A.S. was supported by the European Union’s Horizon 2020 research and innovation program under Marie Skłodowska-Curie grant agreements No 956229 (ALPACA) and No 872539 (PAN-GAIA).

## Disclosure of Interests

The authors have no competing interests to declare that are relevant to the content of this article.

## Footnotes

† It should be noted that the *SNP reconstruction accuracy* can only be computed for SNP data. Therefore this metric should be replaced by another one when using other data types such as gene expression levels.

‡ The hyperparameters of the autoencoder and the PhenoEncoder are equal. This means that the total number of unsupervised learning epochs is equivalent to the total of the number of autoencoder pre-training and the PhenoEncoder training epochs, which is half the joint training time.

§ In this instance, the total number of unsupervised learning epochs for the prototype model is 60, equivalent to the total of autoencoder pre-training (10) and the PhenoEncoder training epochs, which is half the joint training time (100*/*2 = 50). For a fair comparison, the baseline embeddings were obtained via classical, i.e. unsupervised, autoencoder training that lasted 60 epochs.

¶ The baseline latent space is generated by the PhenoEncoder’s multiple autoencoder sub-network, trained for 35 epochs, the sum of pre-training (10) and half the joint training epochs (50*/*2 = 25) on VariantSpark data.

## Notes

https://archive.ics.uci.edu/dataset/342/mice+protein+expression

http://gigadb.org/dataset/100759

https://github.com/gizem-tas/phenoencoder

### Competing Interest Statement

The authors have declared no competing interest.

