## Supplementary Data for "PhenoEncoder: A Discriminative Embedding Approach to Genomic Data Compression"

### Supplementary Note 1

**Autoencoder Building Procedure.** We construct a deep autoencoder model from sequential fully-connected layers, such that the dense connection between consecutive layers can compress the input into a lower dimension at minimum information loss. The following modeling procedure is specifically standardized to comply with varying input sizes. The number of hidden layers $hl$ is set to 4 and corresponds to the number of layers between both the input and output layers and the bottleneck, hence the depths of the encoder as well as the decoder. The total number of layers in the autoencoder then becomes $2 \cdot hl + 3 = 11$.

The number of neurons in each layer $l$ is determined by the hyperparameter *shape*, and is computed using the bottleneck dimension ($bn$), the number of hidden layers ($hl$), and the slope of the encoder and decoder geometries ($p$). $p$ is fixed at 0.5 in this study, corresponding to the *elliptic* shape, see Figure S1. The number of neurons in layer $l$ is then calculated by the function $n(l)$:

$$n(l) = bn + \left( \frac{input\ size - bn}{(hl+1)^p} \cdot |l|^p \right), \ \ where\ l \in \mathbb{Z}, \ -(hl+1) \leq l \leq hl+1. \tag{S1}$$

The number of neurons in the input and output layers are given by $n(-(hl+1))$ and $n(hl+1)$ respectively and are both equal to the input dimension. The number of neurons in a layer should always be a positive integer, otherwise $n(l)$ is rounded to the nearest one. The bottleneck dimension is given by $n(0)$ and is equal to $bn$, a predefined value for a single autoencoder. Alternatively in case of multiple autoencoders, $bn$ is dynamically adjusted based the input size:

$$bn = \min \left( bn_{max}, \frac{input\ size}{10} \right), \ \ where\ bn_{max} = 3 \tag{S2}$$

The weights of the hidden layers are initialized with He uniform variance scaling initializer [41] and we use the Leaky Rectified Linear Unit activation function (Leaky ReLU) in every layer but the output [40].

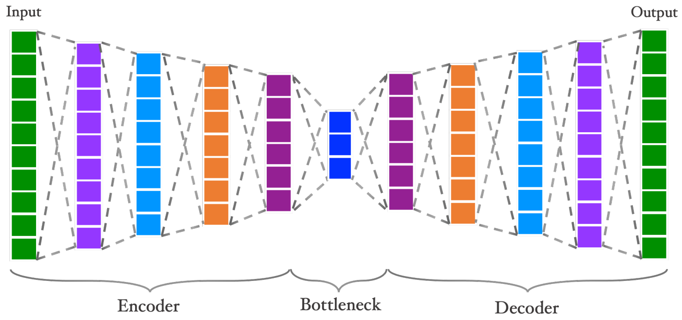


Fig. S1: Visualization of an elliptic autoencoder. This representative model is built with 4 hidden layers and 3 bottleneck nodes for an input of size 10, see also equation (S1).

The output activations depend on the data type. For continuous output values such as protein expression levels, we utilize the linear activation function at the output layer. For genotype data, where the allelic recoding of SNPs ranges from 0 to 2, we fashioned the following custom activation function, $r(x)$, based on the tangent hyperbolic activation function ($tanh(x)$), whose output values are bounded between 0 and 2 in lieu of the original range from -1 to 1:

$$r(x) = tanh(x) + 1 = \left( \frac{e^x - e^{-x}}{e^x + e^{-x}} \right) + 1 \tag{S3}$$

### Supplementary Note 2

**Optimal Hyperparameter Setting.** For the auxiliary classifier, following an unsupervised compression of the original data, we carried out a grid search, using 5-fold cross-validation only on the training samples, concerning the depth (number of hidden layers) along with the width (the number of nodes per layer) of the sub-network, the dropout rate, the learning rate and the batch size.

The grid search for the classifier sub-network of the PhenoEncoder carried out on VariantSpark (training) samples resulted in 2 hidden layers with respective dimensions [32, 8] and 0.1 dropout rate in between, while the optimized prototype of the PhenoEncoder experimented on Mice Protein Expression (training) samples comprised 2 hidden layers with [16, 4] nodes and 0.2 dropout rate.

For the VariantSpark experiment, the optimal number of pre-training epochs was 10 for the autoencoder and 5 for the classifier and the PhenoEncoder's joint training lasted for 50 epochs. As for the prototype model, the autoencoder and the classifier were pre-trained for 10 epochs, while the number of joint training epochs was 100. Finally, all pre-training and joint training steps in both experiments utilized a batch size of 32.

Table S1: Final hyperparameter settings for the PhenoEncoder Experiments.

| **Hyperparameters** | **Prototype Model on Mice Protein Expression Data** | **Full PhenoEncoder on VariantSpark Data** |
|---|---|---|
| **Autoencoder** | | |
| Learning rate | 0.001 | 0.0001 |
| Epochs$^{\star}$ | 10 | 10 |
| Batch size | 32 | 32 |
| **Classifier** | | |
| Depth$^{\star\star}$ | 2 | 2 |
| Width$^{\star\star\star}$ | [16, 4] | [32, 8] |
| Dropout rate | 0.2 | 0.1 |
| Learning rate | 0.001 | 0.0001 |
| Epochs$^{\dagger}$ | 10 | 5 |
| Batch size | 32 | 32 |
| **PhenoEncoder** | | |
| Learning rate | 0.001 | 0.0001 |
| Epochs$^{\ddagger}$ | 100 | 50 |
| Batch size | 32 | 32 |

$^{\star}$ Number of autoencoder pre-training epochs.
$^{\star\star}$ Number of hidden layers.
$^{\star\star\star}$ Number of nodes in the hidden layers.
$^{\dagger}$ Number of classifier pre-training epochs.
$^{\ddagger}$ Number of joint training epochs. The PhenoEncoder is trained for half of the epochs, while the classifier is trained during the remaining half.

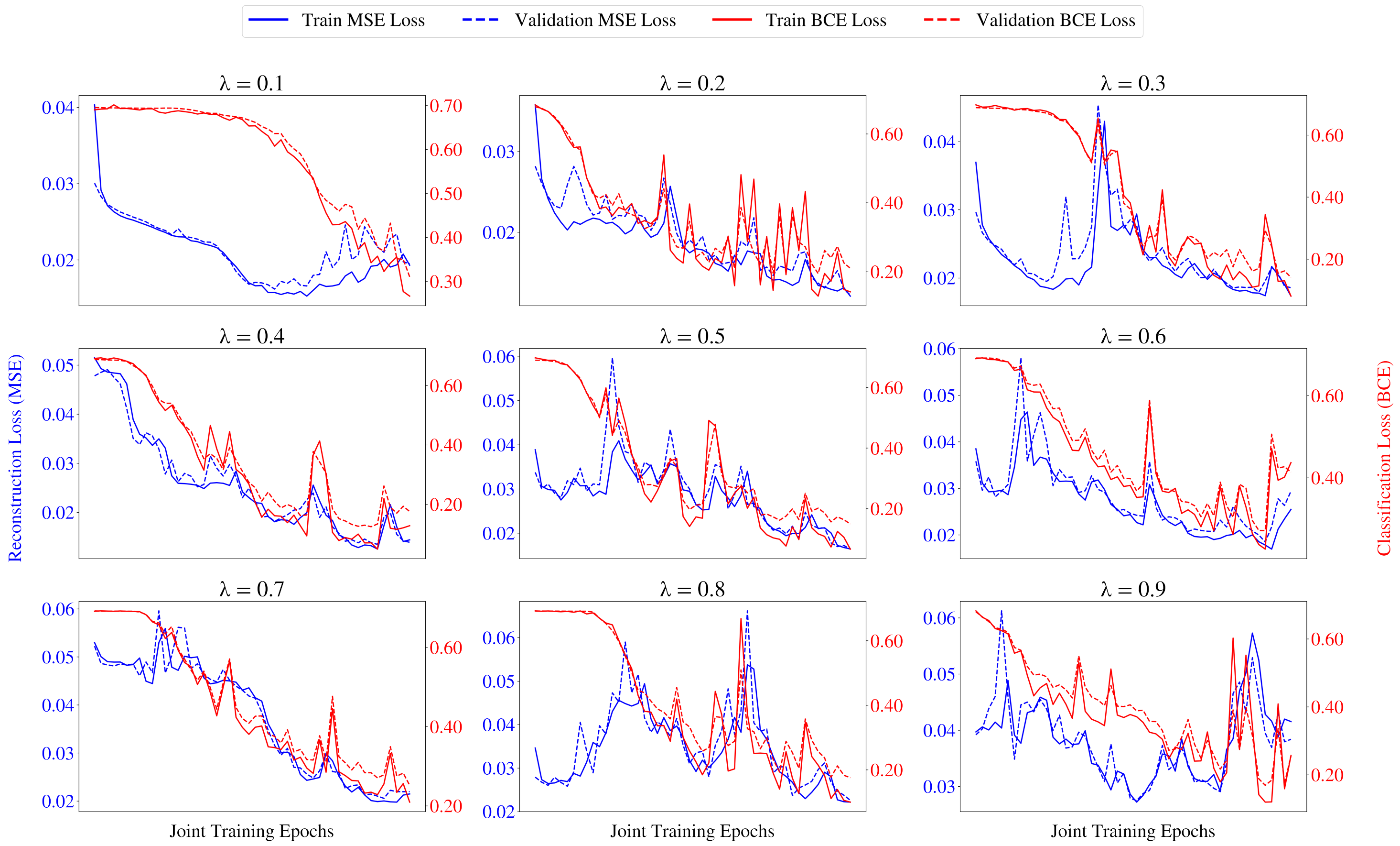


Fig. S2: The progression of the reconstruction (MSE) and classification (BCE) losses throughout their joint optimization compiled under different levels of loss weighting. The models are trained on a randomly selected subset (75%) of the predefined training samples where the remaining 25% constitute the representative validation sample to exemplify the impact of $\lambda$. $\lambda$ represents the relative importance of BCE over MSE in the total loss, therefore defines the delicate balance between them. Smaller values of $\lambda$ could delay the convergence of BCE, while disrupting the progress of an already converged MSE loss. Otherwise for greater $\lambda$, the BCE could dominate the learning until the point of overfitting. Beyond this point, the classifier's loss may unrealistically divert the PhenoEncoder to an invalid latent space (see the fluctuating behaviors of both losses towards the end for 0.8 and 0.9). Then, to ensure a robust feature extraction against noise, the weight of MSE, $1 - \lambda$, matters too.

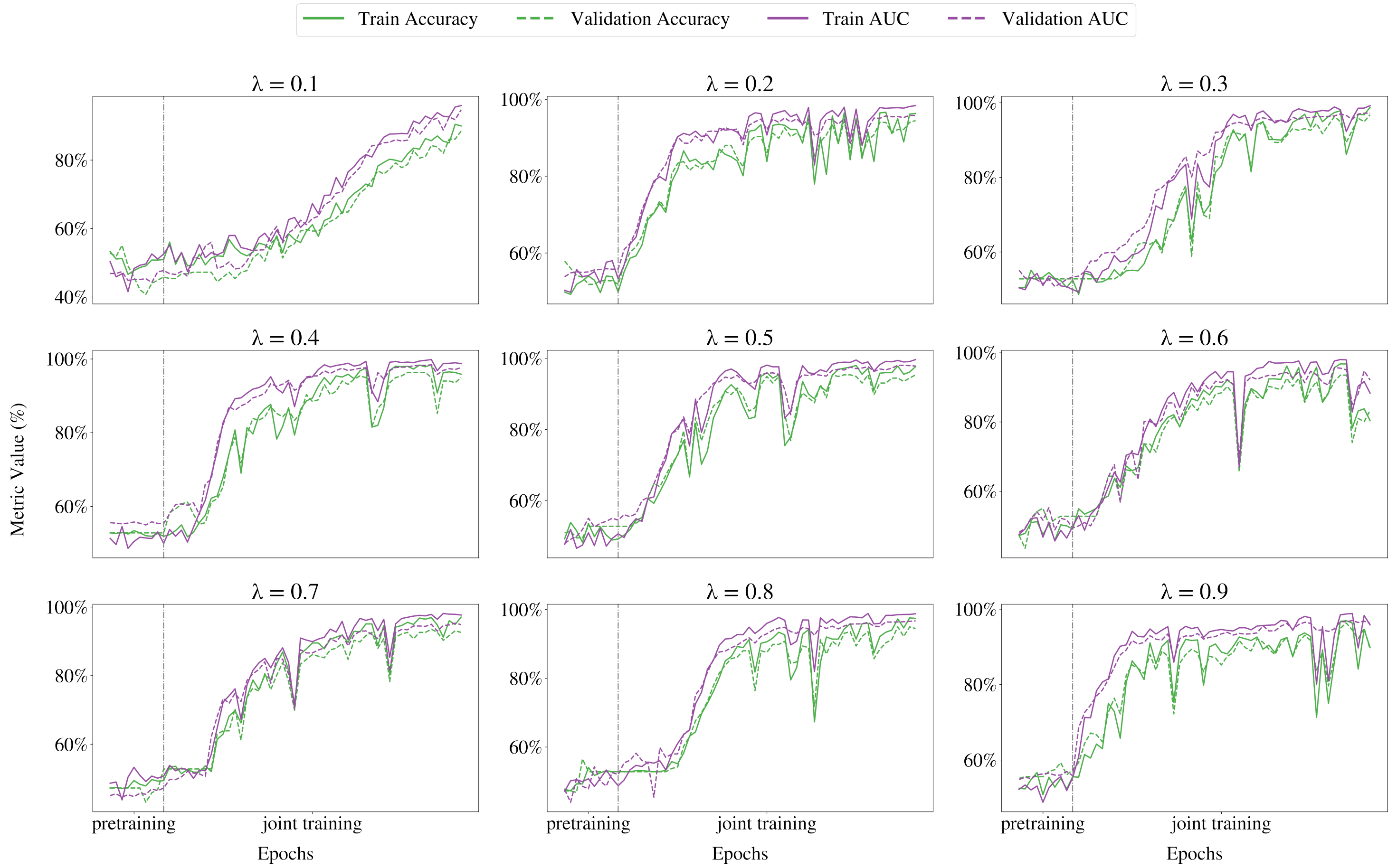


Fig. S3: The impact of different loss weight coefficients on the performance of the auxiliary classifier during joint training. The performance is assessed by means of the classification accuracy and the AUC-ROC metrics. The models are trained on a randomly selected subset (75%) of the predefined training samples where the remaining 25% constitute the exemplary validation sample. While it would be inadequate to deduce an optimal $\lambda$ from these experiments, they still demonstrate that classification performance is not necessarily directly proportional to the weight of the BCE loss, which requires cautious tuning.

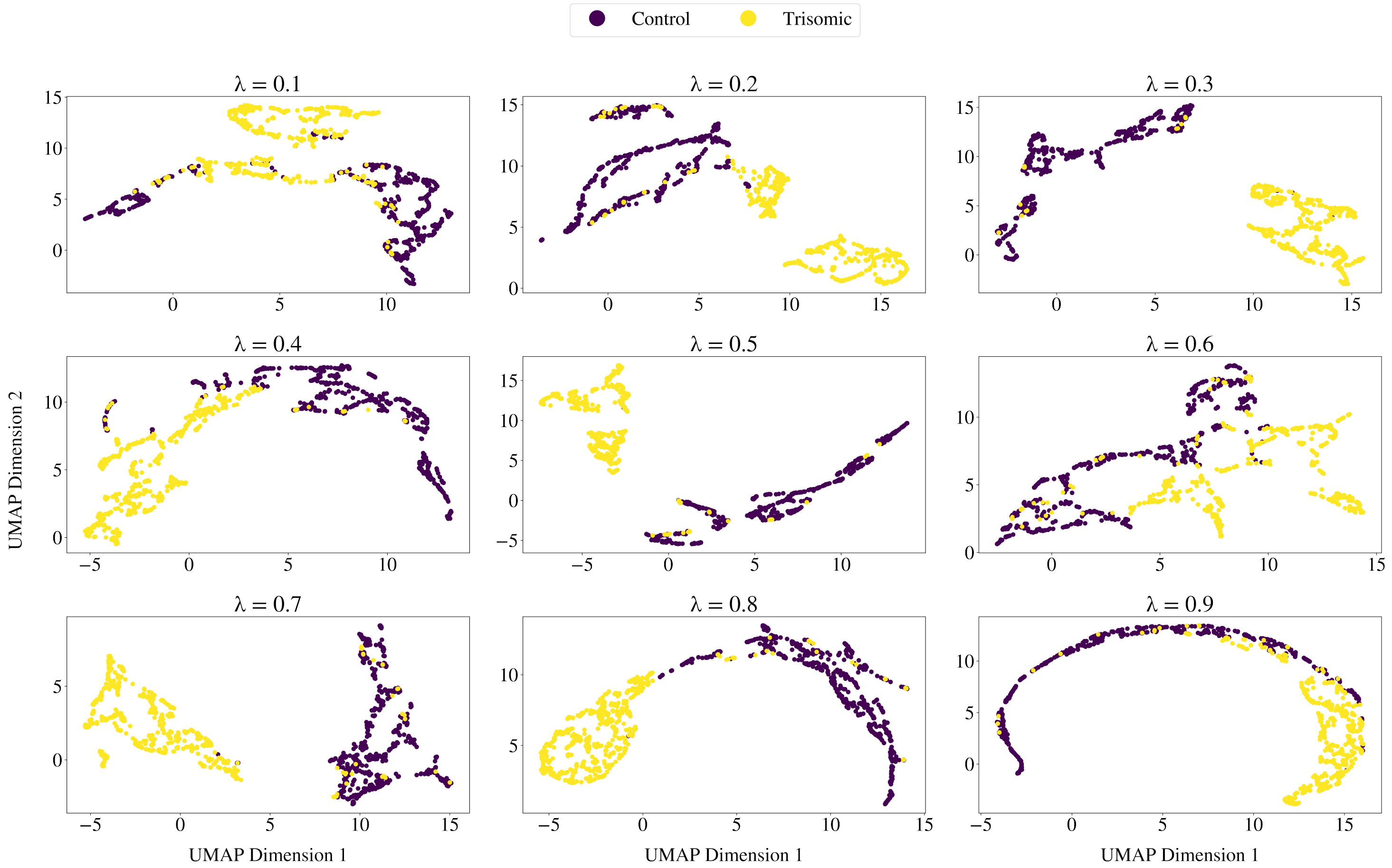


Fig. S4: The impact of different $\lambda$ values on the final embeddings reflected in the visualizations of the latent spaces. The resulting embeddings are projected via UMAP (Uniform Manifold Approximation and Projection) to a 2-dimensional space for practical visualization purposes. The balance between two losses can alter relative positioning of samples in the latent spaces, which reflects the similarity between samples and in fact the learned characteristics of the data. Evidently, $\lambda$ can substantially impact the extent of the discriminative aspects of latent variables which govern the separability of the resulting embeddings.

### Supplementary Note 3

**Loss Weight Coefficient Optimization.** The loss weight coefficient $\lambda$ symbolizes the relative importance of the BCE loss within the joint loss function. A lower $\lambda$ puts emphasis on autoencoder learning, but may reduce the discriminative power of extracted features. Conversely, as $\lambda$ increases, so does the risk of overfitting for the classifier.

To determine the ideal $\lambda$ for this trade-off, we performed 3-fold cross-validation on the training subsets of both datasets, testing $\lambda$ values from 0.1 to 0.9 in increments of 0.1. During this procedure, we stratified the VariantSpark subset by the phenotype and the Mice Protein Expression subset not only by the phenotype but by a composite label that also accounted for behavior and treatment classes, to mitigate spuriousness. For each split, the models were trained using the optimal hyperparameters shown in Table S1, and then evaluated on the corresponding validation fold. The results from the $\lambda$ tuning experiments for the optimal prototype model, listed in Table S2, represent the average metrics across three cross-validation folds per $\lambda$.

Table S2: Cross-validation results for evaluating the prototype model of the PhenoEncoder under different levels of loss weighting. Bold face marks the $\lambda$ value that results in the best performance.

| **Metric** | **Data** | **$\lambda$ value** | | | | | | | | |
|---|---|---|---|---|---|---|---|---|---|---|
| | | 0.1 | 0.2 | 0.3 | 0.4 | 0.5 | 0.6 | **0.7** | 0.8 | 0.9 |
| MSE Loss | Train | 0.0151 | 0.0180 | 0.0233 | 0.0244 | 0.0215 | 0.0222 | 0.0221 | 0.0291 | 0.0434 |
| | Val. | **0.0159** | 0.0180 | 0.0237 | 0.0255 | 0.0222 | 0.0230 | 0.0233 | 0.0300 | 0.0442 |
| BCE Loss | Train | 0.3101 | 0.2396 | 0.1801 | 0.1231 | 0.1562 | 0.1009 | 0.0602 | 0.1600 | 0.2497 |
| | Val. | 0.3369 | 0.3040 | 0.2241 | 0.1963 | 0.1900 | 0.1559 | **0.1315** | 0.1811 | 0.3158 |
| Accuracy | Train | 0.8403 | 0.9178 | 0.9670 | 0.9462 | 0.9618 | 0.9693 | 0.9821 | 0.9537 | 0.9259 |
| | Val. | 0.8264 | 0.8935 | 0.9491 | 0.9248 | 0.9468 | 0.9502 | **0.9583** | 0.9479 | 0.9028 |
| Precision | Train | 0.8625 | 0.9308 | 0.9661 | 0.9912 | 0.9541 | 0.9817 | 0.9976 | 0.9766 | 0.9018 |
| | Val. | 0.8398 | 0.9083 | 0.9461 | **0.9810** | 0.9311 | 0.9638 | 0.9799 | 0.9731 | 0.8793 |
| Recall | Train | 0.7414 | 0.8922 | 0.9645 | 0.8946 | 0.9669 | 0.9534 | 0.9645 | 0.9252 | 0.9571 |
| | Val. | 0.7328 | 0.8627 | 0.9461 | 0.8603 | **0.9583** | 0.9314 | 0.9314 | 0.9167 | 0.9363 |
| AUC-ROC | Train | 0.8966 | 0.9606 | 0.9743 | 0.9947 | 0.9816 | 0.9847 | 0.9958 | 0.9787 | 0.9786 |
| | Val. | 0.8804 | 0.9439 | 0.9703 | 0.9841 | 0.9755 | 0.9758 | **0.9868** | 0.9760 | 0.9630 |
| AUC-PRC | Train | 0.8703 | 0.9625 | 0.9732 | 0.9955 | 0.9669 | 0.9896 | 0.9963 | 0.9843 | 0.9825 |
| | Val. | 0.8560 | 0.9463 | 0.9698 | 0.9861 | 0.9448 | 0.9790 | **0.9884** | 0.9829 | 0.9657 |

We had hypothesized that as the weight of the classification loss increases (as $\lambda$ grows), the autoencoder learning would be gradually compromised. The weight of the MSE loss is the highest for the smallest $\lambda$ ($\lambda = 0.1$). Evidently this is indeed the point where the autoencoders yield the lowest average validation MSE score: 0.0159 (see Table S2). It is also noteworthy that this score does not increase monotonically along the range of $\lambda$, given the slight decrease between 0.4 and 0.5, the junction where both losses become equally weighted.

Given that the phenotype classes are fairly balanced in this dataset, we primarily base the decision-making for optimal $\lambda$ on the overall ability of the model to distinguish between the case and control samples. At $\lambda = 0.7$, the auxiliary classifier yields the lowest average validation BCE loss as well as the highest accuracy, AUC-ROC and AUC-PRC. Figures S5 and S6 indicate elbows at the optimal loss weight of 0.7, beyond which the classifier's performance deteriorates despite the growing emphasis on BCE in the joint loss.

The results from the $\lambda$ tuning experiments for the full PhenoEncoder, shown in Table S3, represent the average metrics across three cross-validation folds for each $\lambda$. As precedented by the prototype model, the best reconstruction performance occurs at $\lambda = 0.1$, indicated by the lowest MSE Loss (0.1260) and the highest SNP accuracy (0.8559). In this case, the MSE increases monotonically (on both training and validation folds), accompanied by a parallel decline in SNP accuracy along the range of $\lambda$.

Table S3: Cross-validation results for evaluating the PhenoEncoder model under different levels of loss weighting. Bold face marks the $\lambda$ value that results in the best performance.

| **Metric** | **Data** | $\lambda$ **value** | | | | | | | | |
|---|---|---|---|---|---|---|---|---|---|---|
| | | 0.1 | 0.2 | **0.3** | 0.4 | 0.5 | 0.6 | **0.7** | 0.8 | 0.9 |
| MSE Loss | Train | 0.1249 | 0.1348 | 0.1396 | 0.1449 | 0.1488 | 0.1501 | 0.1607 | 0.1655 | 0.1707 |
| | Val. | **0.1260** | 0.1357 | 0.1405 | 0.1456 | 0.1495 | 0.1510 | 0.1609 | 0.1663 | 0.1712 |
| BCE Loss | Train | 0.4214 | 0.4170 | 0.3877 | 0.3832 | 0.3858 | 0.3662 | 0.3562 | 0.3110 | 0.3293 |
| | Val. | 0.7579 | 0.7494 | **0.7412** | 0.7684 | 0.7682 | 0.7756 | 0.7616 | 0.8124 | 0.7703 |
| SNP Acc. | Train | 0.8572 | 0.8458 | 0.8405 | 0.8336 | 0.8296 | 0.8263 | 0.8151 | 0.8090 | 0.8027 |
| | Val. | **0.8559** | 0.8448 | 0.8395 | 0.8328 | 0.8287 | 0.8254 | 0.8147 | 0.8083 | 0.8020 |
| Accuracy | Train | 0.8471 | 0.8524 | 0.8824 | 0.8844 | 0.8881 | 0.8924 | 0.8782 | 0.9099 | 0.9026 |
| | Val. | 0.5442 | 0.5499 | 0.5575 | 0.5513 | 0.5224 | 0.5437 | **0.5655** | 0.5482 | 0.5632 |
| Precision | Train | 0.8627 | 0.8394 | 0.8873 | 0.9010 | 0.8702 | 0.8848 | 0.8871 | 0.9144 | 0.8799 |
| | Val. | 0.5463 | 0.5445 | 0.5582 | 0.5523 | 0.5218 | 0.5435 | **0.5658** | 0.5505 | 0.5559 |
| Recall | Train | 0.8268 | 0.8712 | 0.8761 | 0.8637 | 0.9148 | 0.9032 | 0.8686 | 0.9050 | 0.9343 |
| | Val. | 0.5248 | 0.6102 | 0.5498 | 0.5435 | 0.5195 | 0.5435 | 0.5666 | 0.5417 | **0.6466** |
| AUC-ROC | Train | 0.9272 | 0.9312 | 0.9520 | 0.9512 | 0.9561 | 0.9582 | 0.9608 | 0.9722 | 0.9714 |
| | Val. | 0.5737 | 0.5754 | 0.5763 | 0.5690 | 0.5398 | 0.5594 | 0.5795 | 0.5682 | **0.5914** |
| AUC-PRC | Train | 0.9329 | 0.9368 | 0.9523 | 0.9554 | 0.9505 | 0.9565 | 0.9631 | 0.9727 | 0.9711 |
| | Val. | 0.5601 | 0.5702 | 0.5743 | 0.5561 | 0.5415 | 0.5568 | 0.5641 | 0.5659 | **0.5822** |

The values where the lowest validation BCE loss (0.7412 at $\lambda = 0.3$) and the highest accuracy (0.5655 at $\lambda = 0.7$) occur do not align, also illustrated in Figures S7 and S8, nor does the best AUC-ROC (0.5914 at $\lambda = 0.9$). However, the significant decline in reconstruction quality as $\lambda$ increases—given more than 5% drop in SNP accuracy between two extremes—warrants caution. Hence, we omit $\lambda = 0.9$ and proceed with $\lambda = 0.3$ and $\lambda = 0.7$ to the next phase.

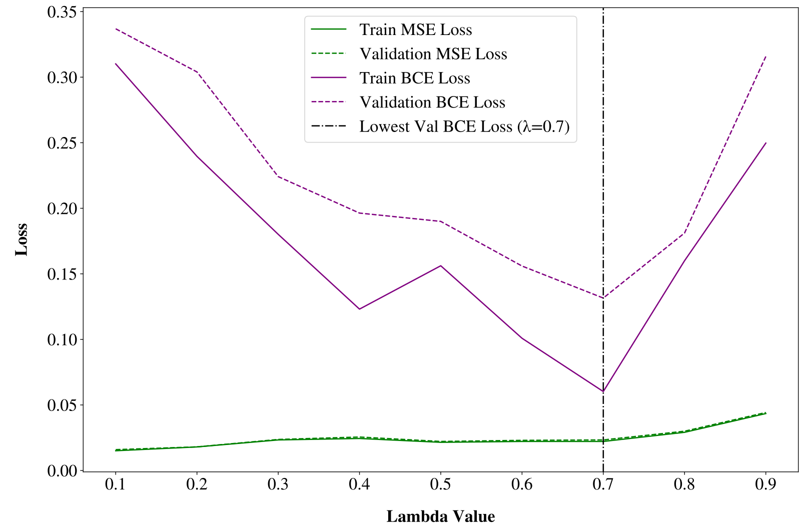


Fig. S5: The train and validation MSE and BCE losses averaged over three cross-validation folds to evaluate the prototype model under nine levels of loss weighting. The black line indicates the lowest BCE loss, hence the best average classification performance at $\lambda = 0.7$, which is the optimal loss weight coefficient.

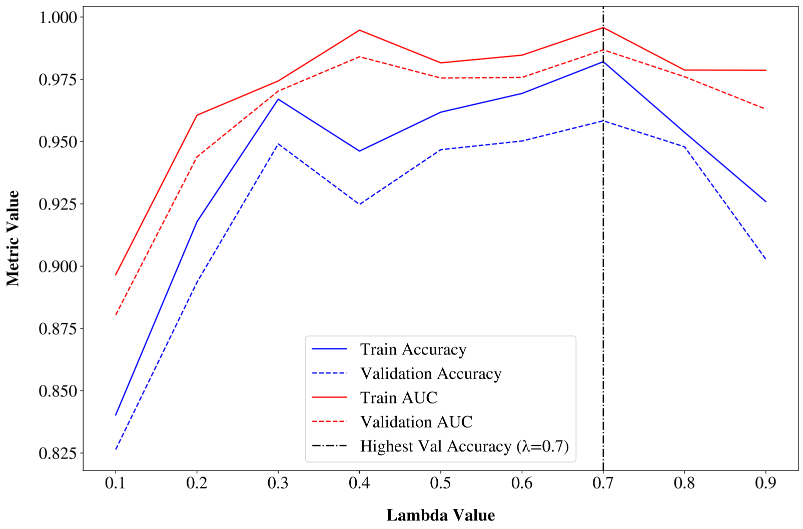


Fig. S6: The train and validation classification accuracy and AUC-ROC metrics, averaged across three cross-validation folds, to evaluate the prototype model under nine levels of loss weighting. The black line highlights the highest average validation accuracy and AUC-ROC, marking the upward elbow at $\lambda = 0.7$.

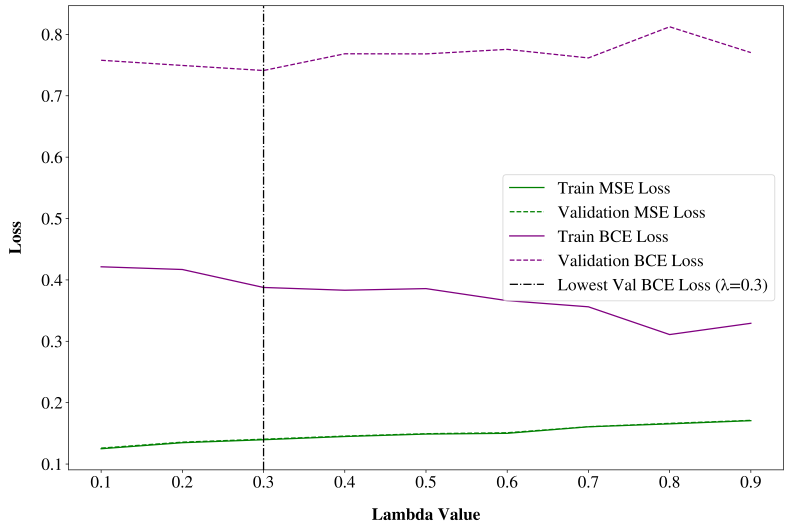


Fig. S7: The train and validation MSE and BCE losses, averaged over three cross-validation folds, to evaluate the full PhenoEncoder under different $\lambda$ values. The black line marks the lowest average validation BCE loss at $\lambda = 0.3$, while the MSE losses show a steadily increasing trend across the x-axis.

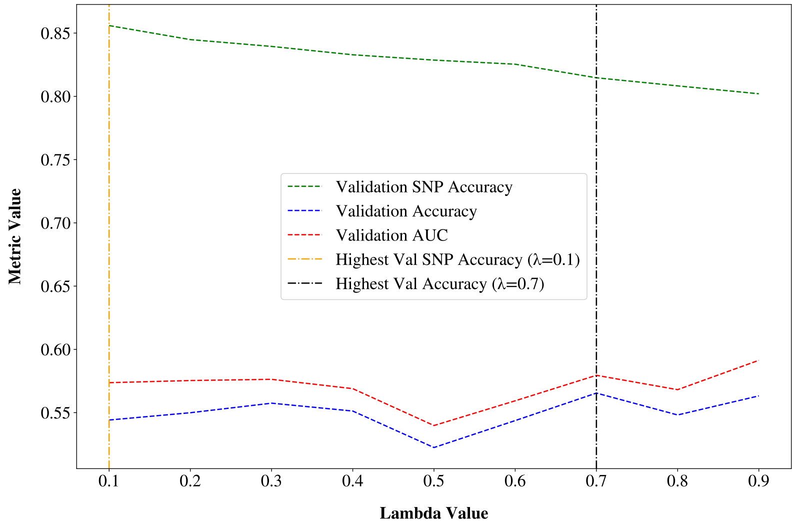


Fig. S8: The validation SNP accuracy, classification accuracy and AUC-ROC metrics, averaged across three cross-validation folds, to evaluate the full PhenoEncoder under different $\lambda$ values. The black line indicates the highest average validation accuracy at $\lambda = 0.7$ with the upward elbow, while the highest validation SNP accuracy is pointed out by the orange line at the smallest $\lambda$ (0.1).

Table S4: Summary of computational resources utilized for the optimization and training of the PhenoEncoder (PE) and the standard autoencoder (AE), as well as the downstream comparison of their feature spaces.

| Task | Node | Cores | CPU Utilized(*) | CPU Efficiency | Job Wall-clock Time(**) | Memory Utilized | Memory Efficiency |
|---|---|---|---|---|---|---|---|
| Classifier Grid Search | CPU | 32 | 08:04:48 | 87.15% | 00:17:23 | 16.74 GB | 13.08% of 128.00 GB |
| $\lambda$ Search | GPU | 2 | 07:26:44 | 55.41% | 06:43:09 | 20.90 GB | 52.26% of 40.00 GB |
| AE Compression | GPU | 2 | 00:23:52 | 62.21% | 00:19:11 | 9.64 GB | 32.15% of 30.00 GB |
| PE Compression | GPU | 2 | 00:37:58 | 58.35% | 00:32:32 | 21.48 GB | 53.71% of 40.00 GB |
| AE vs. PE Comparison | GPU | 16 | 00:10:49 | 7.65% | 00:08:50 | 7.98 GB | 79.76% of 10.00 GB |

*Notes:*

- Time durations are presented in dd-hh:mm:ss format.
- (*): The *CPU Utilized* period indicates the accumulated CPU time across all cores during the job's execution.
- (**): The *Job Wall-clock time* represents the actual elapsed time from the start to the completion of the job in real-time.
